# Individual Variability in the Structural Connectivity Architecture of the Human Brain

**DOI:** 10.1101/2023.01.11.523683

**Authors:** Weijie Huang, Haojie Chen, Zhenzhao Liu, Xinyi Dong, Guozheng Feng, GuangFang Liu, GuoLing Ma, Zhanjun Zhang, Li Su, Ni Shu

## Abstract

The human brain shows higher variability in functional connectivity in the heteromodal association cortex but lower variability in the unimodal cortex. As the anatomical substrate of functional connectivity, the temporal-spatial pattern of individual variability in structural connectivity remains largely unknown. In the present study, we depicted the temporal-spatial pattern of individual variability in structural connectivity, which is highest in the limbic regions and lowest in the unimodal sensorimotor regions. With increasing age, the variability in structural connectivity increased. Our results demonstrated that this specific spatial distribution relates to the plasticity of synapses and white matter. We proposed a modified ridge regression model to predict cognition individually and generate idiographic brain mapping. Individual variability in structural connectivity is significantly correlated with idiographic brain mapping. Overall, our study has potential implications for understanding biological and genetic mechanisms of variability in structural connectivity, guiding interventions to promote successful cognitive ageing and interpreting statistical maps in the human connectome.

## Introduction

Humans think and behave differently from one another, which originates from interindividual variability in cerebral anatomy and function. It is of great importance to understand the neural underpinnings of human identity in neuroscience. The recently released high-quality multimodal neuroimaging datasets, such as the Human Connectome Project ^[1]^ (HCP) and UK Biobank ^[2]^, provide an unprecedented opportunity to investigate neural uniqueness. However, a plethora of evidence suggests that variation in the human brain is not uniform across the brain. For example, long association fibres, e.g., the superior longitudinal fascicle and superior and inferior occipito-frontal fascicles, are more variable than the corticospinal tract, corpus callosum and optic tract in the position and extent of their fibre tracts ^[3]^. In terms of cortical morphology, individual variability in cortical folding is higher in association cortex areas than in motor cortex ^[4]^.

The concept of the human brain connectome plays an important role in comprehending the organization of the human brain. There are two major methods to map human brain networks. One is using diffusing MRI and tractography to construct white matter networks by reconstructing the fibres between areas, and the other is using functional MRI to construct functional networks by measuring interregional neural synchronization. Functional networks act as an identifying fingerprint ^[5]^ and can be used to predict interindividual variation in behaviour ^[6–9]^. There is growing interest in the individual variability in functional connectivity (IVFC). Previous studies have revealed a stable pattern of such variability, with heteromodal association cortices’ variability being significantly higher than that of unimodal cortices ^[10]^. This pattern initially emerges in the third trimester resulting from heterogeneous changes in individual variability across different brain systems, with substantial decreases in the sensorimotor system; intermediate decreases in the visual, dorsal attention, ventral attention and subcortical systems; and slight decreases in the default mode, frontoparietal and limbic systems ^[11]^. In contrast to such decreases in development, IVFC increases with age in adults ^[12]^. The potential causes of the spatial distribution of IVFC remain unclear. A study found that IVFC is positively associated with evolutionary cortical expansion, which is partially retraced in individual development ^[10]^. Therefore, the heterogeneity of IVFC across the cortex may be attributed to the different maturation trajectories. More variable areas, such as the association cortex, are probably less genetically influenced because of prolonged exposure to the environment during maturation. However, a twin study found a complex but significant relationship between genes and IVFC that is age-dependent and bidirectional through systematic voxelwise analyses ^[13]^. In addition, IVFC is positively associated with distant connectivity but negatively associated with local connectivity, and this outcome has been consistently reported across studies ^[10–12]^. This finding seems reasonable because unimodal cortices normally display high local connectivity, while heteromodal association areas show high distant connectivity ^[14]^.

The structural network derived from diffusion magnetic resonance imaging is the anatomical substrate of the functional network. Nevertheless, studies on individual variability in structural connectivity (IVSC) are scarce and have reported inconsistent results. Two studies using the same public dataset, HCP, showed a similar pattern of IVSC with the inferior parietal, temporal, lateral prefrontal cortex and precuneus showing high variability ^[15,16]^. However, another study using a different dataset reported a different spatial distribution of IVSC in which higher variability was localized near language areas, as well as in the medial frontal lobe and in the superior frontal gyrus ^[17]^. The heterogeneous findings may result from several factors: (a) the heterogeneity of the population across studies and (b) methodological differences in the MRI acquisition parameters and methods used to construct structural networks. In addition, these studies all have some deficiencies in IVSC evaluation. For example, the size of the dataset in one study that only included nine participants was too small to obtain a reliable estimation of IVSC. Although another two studies using HCP had large datasets, they included twin subjects, which may lead to IVSC underestimation ^[13]^. None of these studies systematically explored the potential causes and genetic mechanisms of IVSC.

Moreover, machine learning methods have been widely applied to neuroscience research to perform individual-level predictions of human behaviour and cognition ^[18]^. However, traditional machine learning models applying the same features and features’ weights to different individuals are still a population-level predictive approach that relies heavily on multivariate pattern information that is well conserved across individuals. Emerging evidence shows that the functional mapping of the cerebral cortex is variable across individuals ^[19,20]^. Therefore, personalized mapping machine learning methods are particularly important for understanding the neural underpinnings of cognition, as they can capture the complexity of the neural mechanisms and representations of cognition for each person.

The present study encompasses three main objectives. First, we aimed to characterize a reliable and replicable temporal-spatial pattern of IVSC by utilizing multisite datasets. Second, we sought to explore the biological and genetic mechanisms of IVSC by integrating multiomics data. The third objective was to design a machine learning method capable of generating idiographic brain mapping to predict cognition and verify whether the functional mapping of structure connectivity is individualized.

## Results

### Temporal-spatial pattern of IVSC

To delineate the temporal spatial pattern of IVSC, 1724 participants from the HCP Young Adults (HCP-YA) ^[1]^, HCP Retest ^[1]^, HCP Ageing (HCP-A) ^[21]^, Cambridge Centre for Ageing and Neuroscience (Cam-CAN) ^[22]^ and Beijing Ageing Brain Rejuvenation Initiative (BABRI) ^[23]^ were studied. Participants’ demographic information is shown in **Table 1**. Trail A and Trail B tests were used to evaluate the attention and executive function of participants, respectively.

The HCP Retest, where participants underwent two MR scans within one year, was used to estimate intraindividual differences in structural connectivity and regional regression coefficients between interindividual and intraindividual differences in structural connectivity. A small intraindividual difference in structural connectivity indicated high reliability of structural connectivity (**Figure S1**). The intraindividual difference and regression coefficients were applied to the other datasets to estimate IVSC (see Methods for details).

The IVSC of young adults (**Figure 1A**) from the HCP-YA was similar to that of middle-aged and old adults from the combined dataset of HCP-A, Cam-CAN and BABRI (**Figure 1B**) (Spearman’s r = 0.808, p < 0.001). Both IVSC maps showed higher variability in the middle frontal cortex, inferior parietal cortex, middle and inferior temporal cortices, medial frontal cortex, medial temporal cortex and anterior cingulate cortex. To evaluate whether IVSC maps in middle-aged and old adults were stable in different datasets, we harmonized structural networks from the HCP-A, Cam-CAN and BABRI with ComBat ^[24]^ and calculated IVSC maps for the three datasets, respectively (**Figure S2A**, **S2B** and **S2C**). The results showed that IVSC maps in middle-aged and old adults derived from different datasets were not only significantly associated with each other but were also related to the IVSC map derived from the HCP-YA and the combined dataset (**Figure 1C**; all Spearman’s r values > 0.75). The combination of HCP-A, Cam-CAN and BABRI was used for subsequent analyses to explore IVSC patterns across middle-aged and old participants because the estimation of variability is more stable in a larger number of samples. To evaluate whether IVSC is stable over the age range, we split the combined dataset into age tertiles and calculated IVSC for each age group. The 3 IVSC maps were highly similar to each other (**Figure S3**; all Spearman’s r values ≥ 0.97). To assess how the selection of the edge threshold influence the measurement of IVSC, we calculated IVSC maps on the combined dataset after removing weak edges under different edge thresholds. The IVSC maps after removing weak edges were highly similar to the IVSC map without removing edges (**Figure S4**; all Spearman’s r values > 0.96). Additionally, we found that IVSC significantly increased with age in almost all brain regions by using sliding window analysis (**Figure 1D**). Additionally, the mean IVSC and standard deviation of IVSC were both positively correlated with age (**Figure 1E** and **1F**).

**Figure 1.**
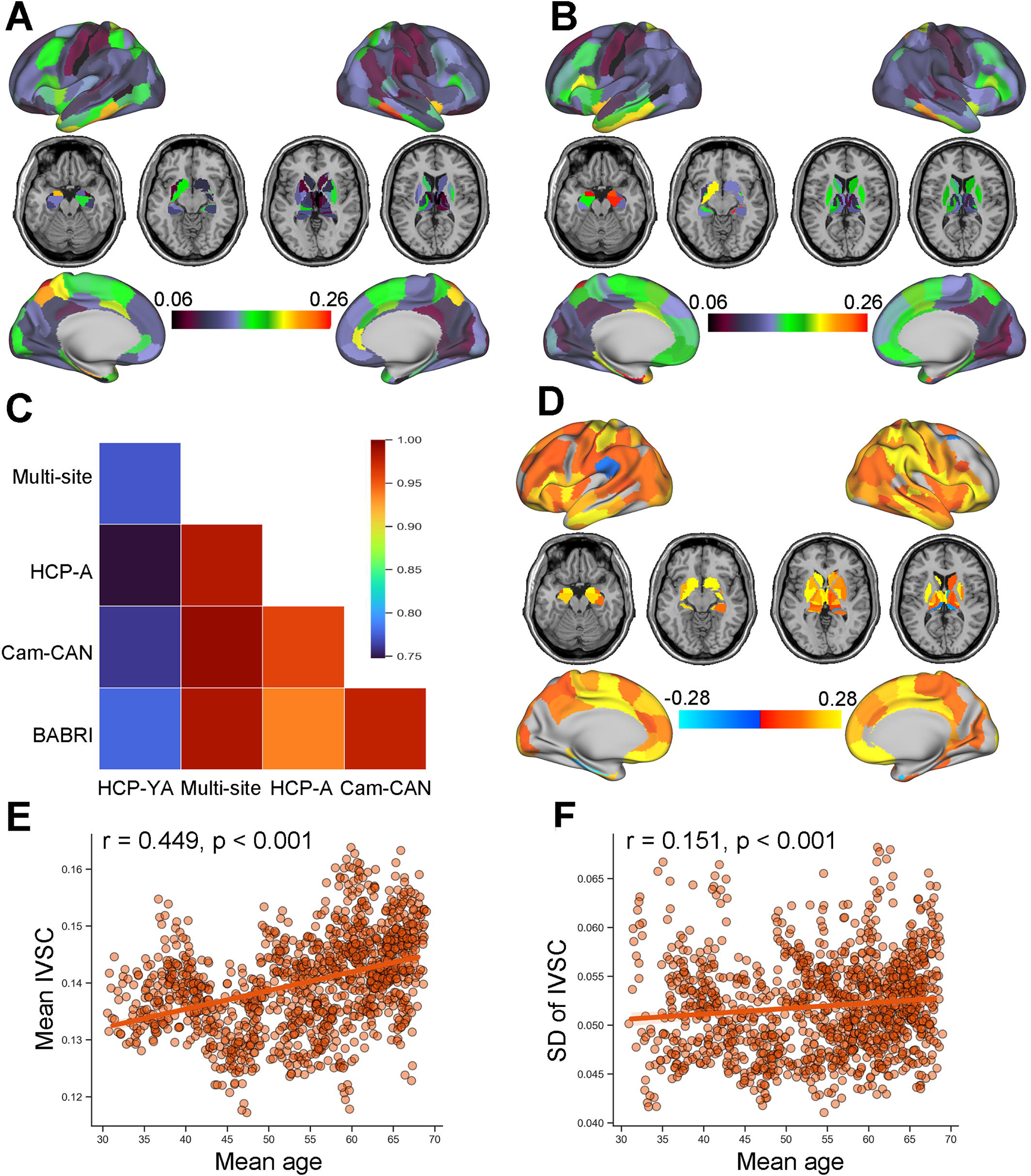
Temporal spatial pattern of IVSC. **A**, The IVSC across participants in HCP-YA. **B**, The IVSC across participants in the combination of HCP-A, Cam-CAN and BABRI studies. **C**, The correlation between IVSC values from different datasets. **D**, Age-related rates of IVSC obtained by performing sliding window analysis in the combined dataset of HCP-A, Cam-CAN and BABRI. **E**, The relationship between mean IVSC and mean age. **F**, The relationship between the standard deviation of IVSC and mean age. IVSC, individual variability in structural connectivity; HCP-YA, Human Connectome Project Young Adults; HCP-A, Human Connectome Project Ageing, Cam-CAN, Cambridge Centre for Ageing and Neuroscience; BABRI, Beijing Ageing Brain Rejuvenation Initiative; SD, standard deviation.

IVSC was also assessed within 8 brain systems, including 7 cortical networks ^[27]^ (**Figure 2A**) and a subcortical system. The limbic network had the highest level of IVSC, whereas the somatomotor network and visual network were least variable (**Figure 2B**). A sliding window analysis revealed that mean variability in all 8 brain systems significantly increased with age (**Figure 2C**).

**Figure 2.**
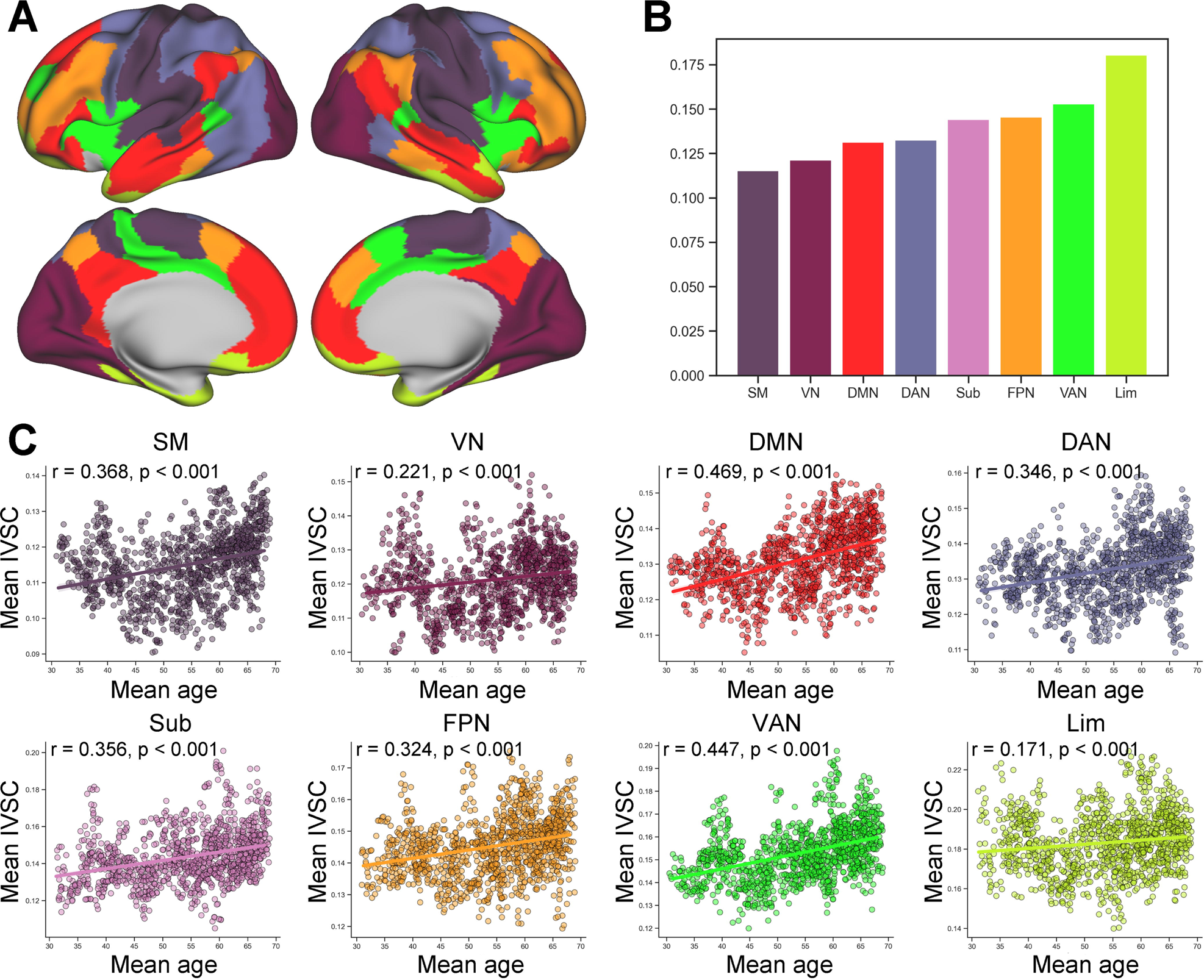
IVSC quantified in 8 brain systems. **A**, The spatial distribution of 7 cortical networks defined by Yeo and his colleagues ^[27]^. **B**, Mean IVSC in 8 brain systems. **C**, The relationship between mean IVSC in 8 brain systems and mean age. IVSC, individual variability in structural connectivity; SM, somatomotor network; VN, visual network; DMN, default mode network; DAN, dorsal attention network; Sub, subcortical system; FPN, frontosparietal network; VAN, ventral attention network; Lim, limbic network.

We also replicated the above analyses with different atlases, Automated Anatomical Labelling (AAL) atlas ^[25]^ and a cortical parcellation created by Gordon and his colleagues ^[26]^. The IVSC maps under different atlases were similar, with all maps showing higher variability in medial prefrontal cortex, medial temporal cortex, anterior cingulate cortex and middle frontal cortex, and lower variability in superior temporal cortex, precuneus, posterior cingulate cortex and the cortex near central sulcus (**Figure S5A** and **S5E**). In addition, the relationship between IVSC and age remained stable (**Figure S5B-D** and **S5F-H**). We mapped AAL atlas into 7 cortical networks and a subcortical system, and mapped Gordon’s atlas into 7 cortical networks because it does not include subcortical regions. The networks with highest or lowest variability are consistent across different atlases, but the order of networks with moderate variability is slightly different across the 3 atlases (**Figures S6A** and **S7A**). The significant increases of mean IVSC in all brain systems were reproduced with using AAL and Gordon’s atlases (**Figure S6B-I** and **S7B-H**).

### Association between IVSC and other properties of brain organization

Having identified the specific temporal spatial pattern of IVSC, we next sought to understand the biological mechanism underlying this specific spatial distribution of IVSC. The human cortex, regardless of the neocortex and allocortex, consists of several layers, which have a characteristic distribution of different neurons and connections with other cortical and subcortical regions. A previous study in mammals demonstrated that the existence of connections is not only associated with physical distance but is also related to the cytology of cortical areas ^[28]^. In addition, myelin proteins can inhibit axon sprouting and stop the formation of structural connections by collapsing the tips of growing axons ^[29]^. Hence, we directly linked IVSC to these fundamental properties of brain organization. Using spatial permutation testing, we found that high IVSC was related to low laminar differentiation (**Figure 3D**; Spearman’s r = −0.461, p = 0.002), low myelin content (**Figure 3E**; Spearman’s r = −0.442, p < 0.001) and low strength of short SC (**Figure 3F**; Spearman’s r = −0.421, p < 0.001). To evaluate the stability of the correlation between IVSC and the strength of short SC, we filtered white matter networks with different edge thresholds. Then, we calculated the IVSC and performed a correlation analysis for each edge threshold. We found that the correlation remained significant under different edge thresholds (**Figure S10**; −0.43 ≤ Spearman’s r ≤ −0.480).

**Figure 3.**
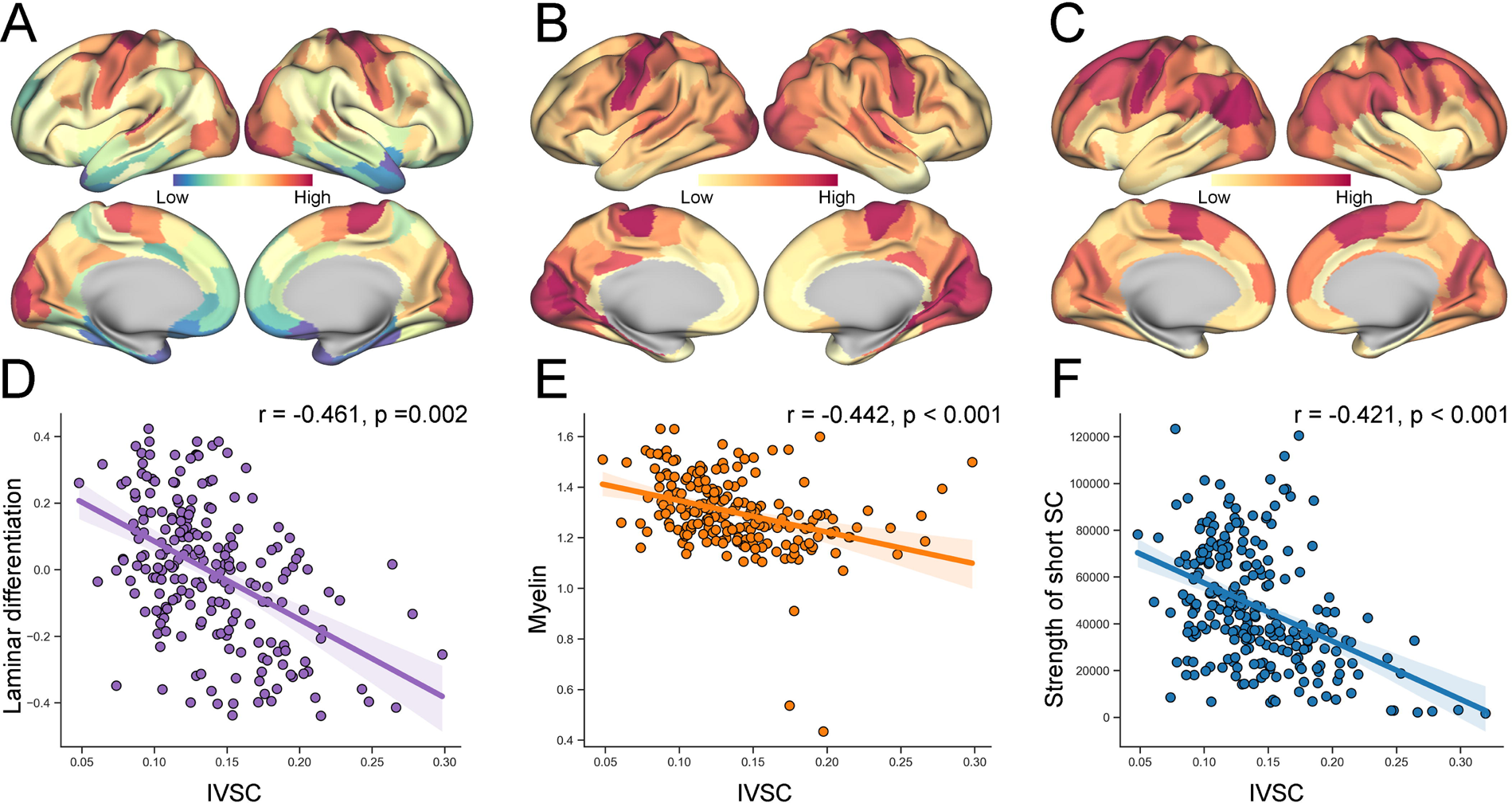
IVSC relates to fundamental properties of brain organization. **A**, Spatial distribution of the laminar differentiation identified by Paquola et al ^[62]^. **B**, Spatial distribution of myelin content measured by T1w/T2w mapping ^[63]^. **C**, Spatial distribution of short connectivity strength. **D-F**, IVSC was associated with the laminar differentiation, myelin content and strength of short connectivity. IVSC, individual variability in structural connectivity; SC, structural connectivity. IVSC, individual variability in structural connectivity; SC, structural connectivity.

For AAL atlas, we only replicated the analysis evaluating the relationship between IVSC and strength of short SC, because it is a volumetric atlas but laminar differentiation and myelin are surface-based measures. For Gordon’s atlas, we re-estimated the above three relationships. Myelin and strength of short SC were consistently significant with IVSC (**Figure S8** and **S9B-C**).

### Association between IVSC and gene expression

Next, we explored the genetic mechanisms of IVSC using a public transcriptional dataset. For the Allen Human Brain Atlas (AHBA), after controlling the quality of gene expression and filtering nonbrain-specific genes, the expression levels of 2112 brain-specific genes across 123 left brain regions were obtained. Using partial least square (PLS) analysis, a multivariate statistical method to extract covarying components from matrices, we found a component of gene expression positively related to IVSC (**Figure 4B**; Spearman’s r = 0.482, p = 0.005) and accounting for 26.1% of the spatial variance in IVSC. The component of gene expression was the sum of weighted gene expression, which we referred to as a gene score. The medial temporal lobe, anterior cingulate cortex and insula showed the highest gene scores, while the precentral gyrus, postcentral gyrus, and occipital gyrus showed the lowest gene scores (**Figure 4A**).

**Figure 4.**
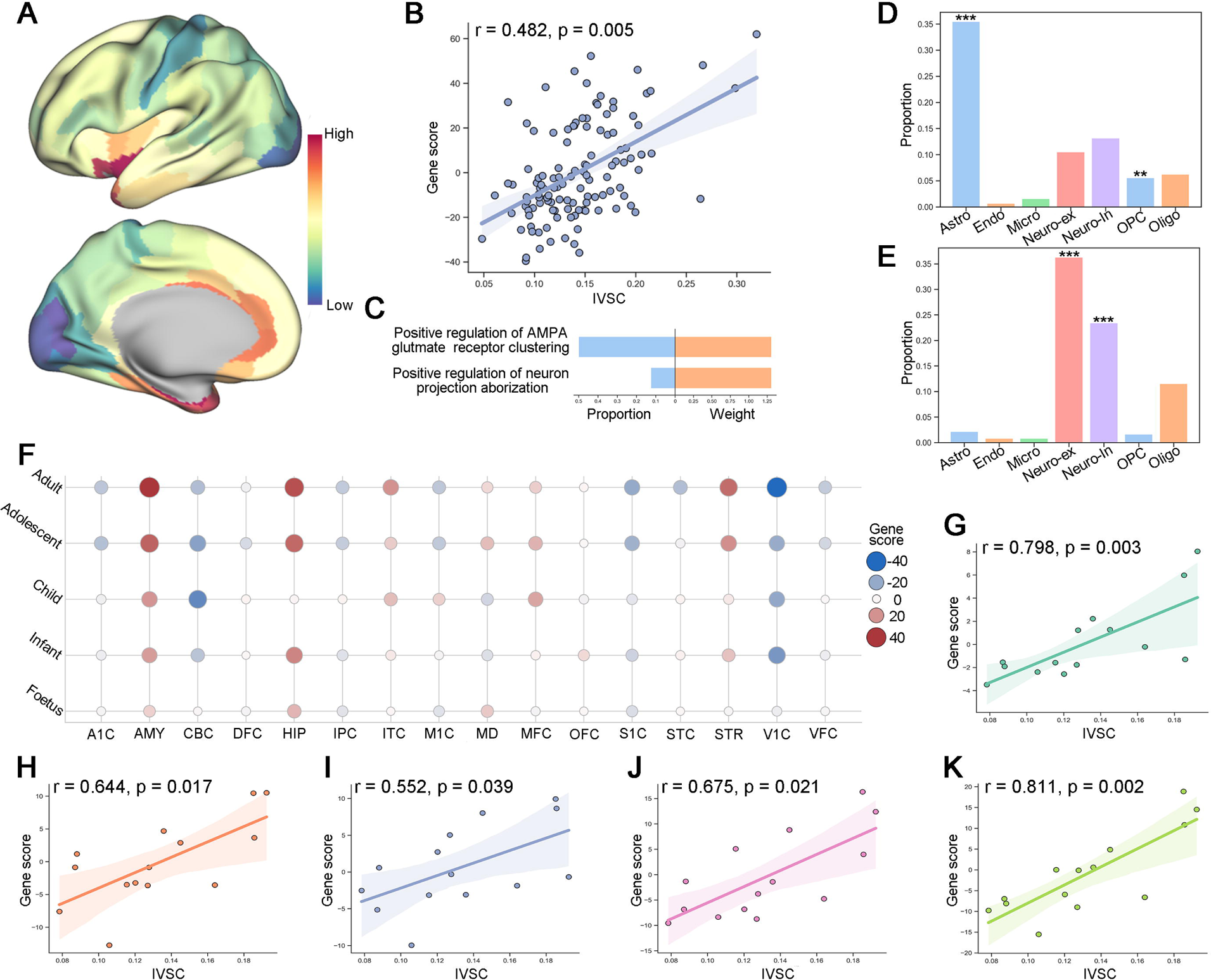
IVSC relates to gene expression profiles. **A**, Spatial distribution of the gene expression score identified by PLS. **B**, IVSC was correlated with the gene expression score. **C**, Biological processes enriched in the positive gene set were identified by performing gene set enrichment analysis and were tested against a spatial autocorrelation-preserving null model. **D**, The proportion of genes in each gene set preferentially expressed in seven different cell types. **F**, Gene scores through human development. **G-K**, IVSC was associated with gene score in foetus, infant, child, adolescent and adult groups. IVSC, individual variability in structural connectivity; Astro, astrocyte; Micro, microglia; OPC, oligodendrocyte precursor; Oligo, oligodendrocyte; Endo, endothelial; Neuro-ex, excitatory neurons; Neuro-in, inhibitory neurons.

Based on the weights derived from the above PLS analysis, we calculated the gene scores of brain regions parcellated by AAL and Gordon’s atlases. For AAL atlas, the gene score is significantly correlated with IVSC (**Figure S11**). For Gordon’s atlas, the gene score is marginally significantly associated with IVSC (**Figure S12**).

The relationship between IVSC and gene score was determined by the genes that contributed most to the gene score. The weights of genes reflected the shared variance between levels of gene expression and gene score. We defined the genes with the top 50% positive/negative weights as strongly contributing genes. To explore the biological processes of the positive and negative gene sets, we performed gene set enrichment analyses in both gene sets, respectively. We identified two biological processes, “positive regulation of AMPA glutamate receptor clustering” (p = 0.049) and “positive regulation of neuron projection arborization” (p = 0.049), in the positive gene set but revealed no biological process in the negative gene set (**Figure 4C**).

Having found the biological process enriched in the positive gene set, we next investigated whether the genes associated with IVSC were preferentially expressed in specific cell types. Specifically, the proportion of genes that were preferentially expressed in each cell type was calculated. Permutation tests were used to assess statistical significance. Positive genes were significantly more highly expressed in astrocytes (p < 0.001) and oligodendrocyte precursors (p = 0.004) (**Figure 4D**), while negative genes were significantly more highly expressed in excitatory (p < 0.001) and inhibitory (p < 0.001) neurons (**Figure 4E**).

In addition, we used the BrainSpan dataset to evaluate the change in gene score throughout human development. The gene score of each tissue sample was calculated based on the gene weights derived from the above PLS analysis. The tissue samples were separated into five life stages (foetus, infant, child, adolescent and adult) in terms of age. The gene scores of tissue samples from the same life stage and same region were averaged to yield a gene score per region and life stage (**Figure 4F**). The gene scores in most regions increased with development. We also used the BrainSpan dataset to evaluate the reproducibility of the association between IVSC and gene scores. We found that IVSC significantly correlated with gene scores in the five life stages (**Figure 4G-K**; foetus: Spearman’s r = 0.798, p = 0.003; infant: Spearman’s r = 0.644, p = 0.017; child: Spearman’s r = 0.522, p = 0.039; adolescent: Spearman’s r = 0.675, p = 0.021; and adult: Spearman’s r = 0.811, p = 0.002). Notably, correlation analyses were performed across 16 brain regions with gene expression levels (see Methods for more details).

### Individualized feature selection improves the prediction accuracy of structural connectivity

Considering that the variability in structural connectivity may arise from plastic changes in white matter, the structural connectivity involved in cognitive tasks may be different across individuals. To verify the hypothesis, we modified the traditional ridge regression to the model that can select features individually to predict cognitions by integrating the rectified linear unit activation function into ridge regression (see Methods for more details), and we investigated whether adding individualized feature selection into the traditional machine learning method can improve the accuracy of cognitive prediction by comparing the performance of traditional and modified ridge regression (**Figure 5A**). The mean absolute error (MAE) and Pearson r value were used to evaluate the models’ performance. Notably, we excluded participants whose scores were less than 10 or more than 300 when predicting executive function and participants whose scores were less than 10 when predicting attention. These participants’ cognitive assessments may not have been reliable.

**Figure 5.**
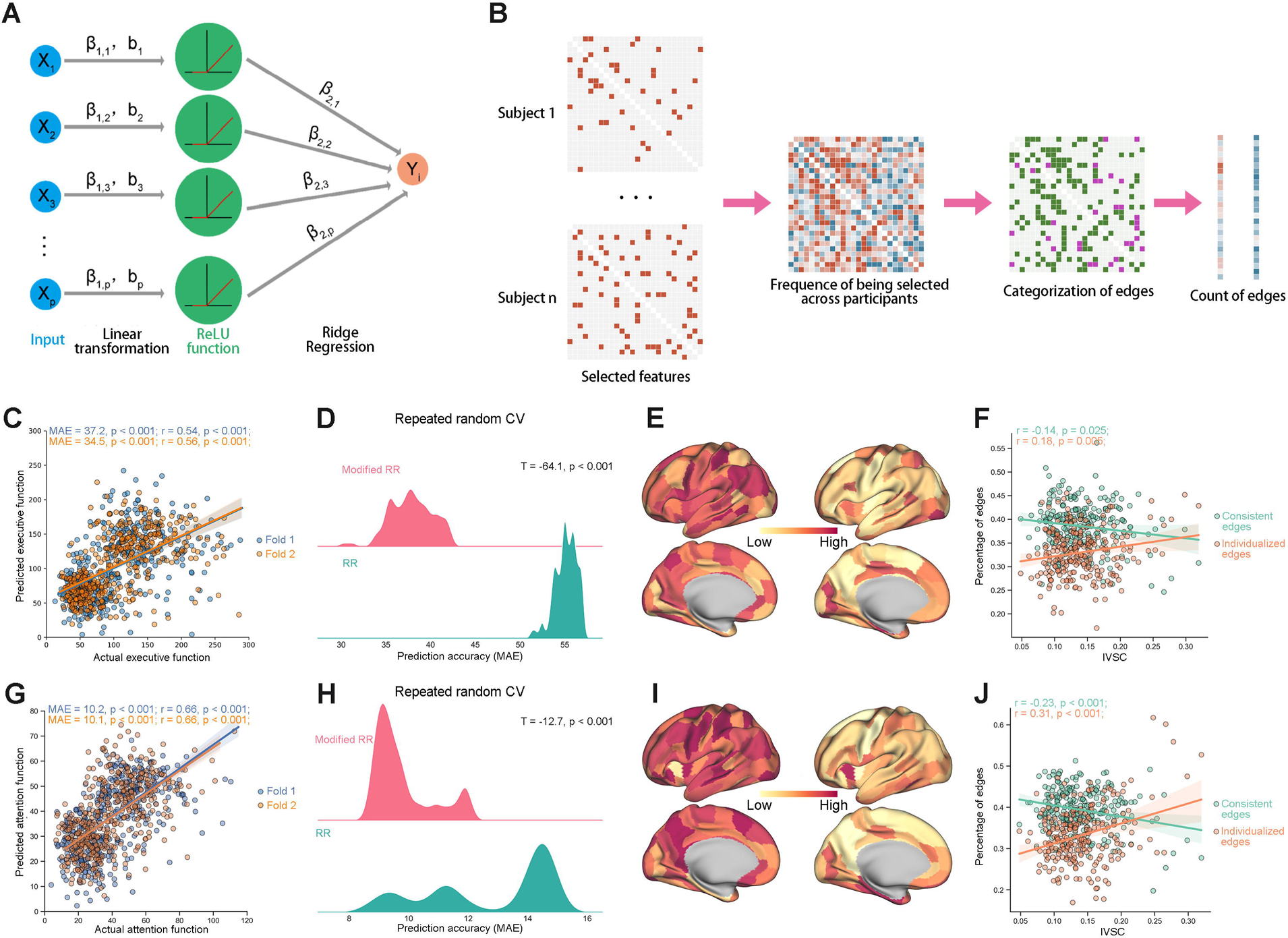
Prediction of executive function and attention benefits from individualized feature selection. **A**, Ridge regression was modified by integrating the ReLU activation function into it. **B** For each edge, the number of participants who selected it as a feature was stated. Then, the edges that were selected by at least two-thirds of participants were defined as consistent edges, and the edges that were selected by less than a third of participants were defined as individualized edges. Finally, we calculated the percentage of consistent edges and individualized edges for each node. **C**, Structural connectivity can be used to predict executive function with modified ridge regression. **D**, Modified RR outperformed RR in the task of predicting executive function. **E**, Percentage of consistent edges (left) and individualized edges (right) in the prediction of executive function. **F**, Percentage of consistent edges and individualized edges related to IVSC across brain regions when predicting executive function. **G**, Structural connectivity can be used to predict attention function with modified ridge regression. **H**, Modified RR outperformed RR in predicting attention function. **I**, Percentage of consistent edges (left) and individualized edges (right) in the prediction of attention. **J**, Percentage of consistent edges and individualized edges related to IVSC across brain regions when predicting attention function. ReLU, rectified linear unit. MAE, mean absolute error. CV, cross validation. RR, ridge regression. IVSC, individual variability in structural connectivity.

Structural connectivity can be used to accurately predict executive function (**Figure 5C**; fold 1: MAE = 37.2, p < 0.001; Pearson’s r = 0.54, p < 0.001; fold 2: MAE = 34.5, p < 0.001; Pearson’s r = 0.56, p < 0.001) and attention (**Figure 5G**; fold 1: MAE = 10.2, p < 0.001; Pearson’s r = 0.66, p < 0.001; fold 2: MAE = 10.1, p < 0.001; Pearson’s r = 0.66, p < 0.001) in matched split-half samples with modified ridge regression. Repeated random 2-fold cross validation (2F-CV) produced similar results when predicting executive function (**Figure 5D**, mean MAE = 37.9, mean Pearson’s r = 0.52) and attention (**Figure 5H**, mean MAE = 9.8, mean Pearson’s r = 0.67) and thus suggested that the matched split of the data was unbiased. We also compared the performance of modified and traditional ridge regression models derived from repeated random 2F-CV by using a two-sample t test and found that the modified ridge regression models outperformed the traditional ridge regression models in both tasks (**Figure 5D**; executive function: T = −64.1, p < 0.001; **Figure 5H**; attention function: T = −12.7, p < 0.001).

For each edge, we counted the participants who selected it as a feature when predicting cognition. Then, we defined an edge that was selected by at least two-thirds of participants as a consistent edge and an edge that was selected by less than one-third of participants as an individualized edge (**Figure 5B**). Finally, we found that the percentage of consistent edges (**Figure 5E**, **I**, left) was negatively associated with IVSC, while the percentage of individualized edges (**Figure 5E, I**, right) was positively associated with IVSC when predicting executive function (**Figure 5F**; consistent edge: Pearson’s r = −0.14, p = 0.025; individualized edge: Pearson’s r = 0.18, p = 0.005) and attention (**Figure 5J**; consistent edge: Pearson’s r = −0.23, p < 0.001; individualized edge: Pearson’s r = 0.31, p < 0.001).

## Discussion

In this study, we leveraged a human connectome method and multicentre data to delineate the spatiotemporal pattern of IVSC and seek to explore the biological mechanism and significance of IVSC distribution. Building upon findings from this study, we identified a robust spatial distribution of IVSC with the limbic network showing the highest variability, association networks showing moderate variability and unimodal networks showing the lowest variability. IVSC increased with ageing in almost all cerebral areas. In addition, we provided evidence that IVSC was strongly linked to the microscopic anatomical properties of the brain and gene expression. Finally, we demonstrated that individualized feature selection improved the prediction accuracy of structural connectivity. Taken together, these results offer a new perspective of brain organization and highlight the importance of individual effects in cognitive neuroscience research.

The spatial distribution of IVSC we observed was moderately different from that in previous studies, although these studies all showed the lowest variability in the visual and somatomotor networks ^[15–17]^. The inconsistent findings may mainly be due to selection bias. The sample size of a previous study was only nine, which was too small to represent the population. In addition, decreased IVFC was associated with increased genetic sharing magnitude ^[13]^. Therefore, including twins in the other two studies may have led to an underestimation of IVSC, especially in regions with high heritability. Compared with FC, the interindividual variability in SC is lower on average than in FC in middle-aged and old adults and exhibits a different spatial pattern mainly in the limbic network (i.e., temporopolar and orbitofrontal area), which showed the highest variability in SC but moderately lower variability in FC. Intuitively, this disassociation is possible because resting-state functional connectivity not only reflects the underlying anatomical circuitry but also recapitulates the history of experience-driven coactivation ^[30,31]^.

Of note, the spatial distribution of IVSC is highly consistent with the sensory-fugal gradient, where one pole is occupied by the limbic areas, which are polarized towards the internal milieu and responsible for the regulation of memory, motivation, emotion and autonomic-endocrine function, and the opposite pole is the primary sensory-motor area, which is polarized towards the extrapersonal environment and provides a gateway for the entry of sensory information into the cerebral neural circuitry and for outputting motor programs to bulbar and spinal motoneurons ^[32]^.This distribution was proposed based on anatomical, physiological and behavioural experiments in macaque monkeys and was demonstrated in the human brain with cytoarchitectonics, functional imaging, electrophysiological recordings and the behavioural effects of focal lesions ^[32]^. The areas further away from the external input have more variable white matter connections across humans. In the deep learning field, the connections in the earlier and middle layers of the architecture neural network are interpreted as a representation of the data and are considered generally applicable across similar tasks, but the connections in the latter layers of the architecture neural network are task specific. Based on the theory, transfer learning, an approach to reuse a pretrained model on a new problem, is widely applied to a variety of scenarios. In transfer learning, the earlier and middle layers remain relatively consistent across different tasks, whereas the latter layers are variable across different tasks. The shared structural characteristics between the human brain network and architecture neural network may suggest that the flexibility of cerebral latter layers, namely, transmodal areas, is sufficient to guarantee that one has the ability to adapt to changes in the environment, and the stability of sensor-motor areas plays a critical role in the accurate registration of new inputs.

Having identified the spatial distribution of IVSC, we sought to understand whether the macroscopic pattern of structural connectivity is constrained by microscopic properties of brain structure. We observed that higher IVSC is associated with better laminar differentiation and less myelin content. Recent evidence has demonstrated that laminar differentiation is related to the cortical plasticity-stability continuum of areas. The areas with worse laminar differentiation have more markers of synaptic plasticity (i.e., glial fibrillary acidic protein) and fewer markers of stability (i.e., parvalbumin), while the areas with better laminar differentiation show the opposite trend ^[33]^. In addition, intracortical myelin is another marker that limits synaptic plasticity because the proteins in myelin, including Nogo-A ^[34,35]^, MAG ^[36]^ and OMgp ^[37,38]^, can lead to the collapse of tips of growing axons and thus prevent projections towards their target. Areas with higher plasticity are more easily influenced by life experience and may be more variable across populations. Hence, the heterogeneous spatial distribution of IVSC may be a macroscopic reflection of the difference in synaptic and axonal plasticity across different brain regions.

The relationship between IVSC and the strength of short connectivity suggests that the variation can be mainly sourced from the plastic change in weak short white matter connectivity. However, this type of weak short connectivity may be noise because of the limitation of the tractography algorithm, although we used the most cutting-edge processing pipeline and evaluated the correlation under different edge thresholds. Further studies are needed to prove this hypothesis. We also found that IVSC increased with ageing, which is consistent with the findings in a study of functional connectivity ^[12]^. The age-related increase in IVSC indicates that the plastic change may last across the lifespan. The potential neuropathology of neurodegenerative disease may contribute to part of the increase in variation, although we only included relatively young and healthy participants.

To further understand the genetic mechanism underlying the distribution of IVSC, we used expression data from AHBA and found two enriched biological processes: “positive regulation of AMPA glutamate receptor clustering” and “positive regulation of neuron projection arborization”. The AMPA receptor, a target of glutamate, has been proven to be important for modifying synaptic efficacy ^[39]^. Therefore, both biological processes are related to changes in neural pathways, which are the anatomical infrastructure for supporting learning. In addition, we observed that areas with higher IVSC are enriched for genetic signals of astrocytes and oligodendrocyte precursors. Interestingly, axonal electrical activity not only mediates the proliferation of oligodendrocyte precursors by controlling the production and release of growth factors ^[40]^ but also triggers the release of cytokine leukaemia inhibitory factor by astrocytes, which promotes myelination ^[41]^. Combining this evidence, the areas with higher IVSC may have higher expression of genes involved in the formation of new pathways and the strengthening of existing pathways. Hence, the differences in synaptic and axonal plasticity across brain regions may result from the selective expression of the related genes.

Moreover, our results demonstrated that IVSC is linked to executive and attention function. We found that the regions with higher IVSC have more variable representations of cognition across individuals. This is in contrast to the findings of traditional neuroimaging studies, in which a statistical map was computed across a group of subjects. Specifically, traditional neuroimaging studies are more likely to obtain a significant result in areas with low variability but less likely to obtain a significant result in areas with high variability. Hence, consideration of individual effects can help improve the present understanding of individual differences in behaviour.

Several methodological considerations in the present study should be noted. First, the structural networks we used to estimate IVSC were constructed based on a population-based atlas. However, the spatial topological properties of cortical regions vary across individuals ^[20,42,43]^. Therefore, using a population-based atlas may be disadvantageous to estimate IVSC. Second, we regressed out intraindividual variability from interindividual variability, but the intraindividual variability and interindividual variability were estimated in different datasets, which may also influence the estimation of IVSC. Third, the transcriptional data that we used were based on small samples of post-mortem brains. More comprehensive microarray gene expression datasets are necessary for future studies.

In summary, we provided evidence that IVSC is constrained by microanatomical properties and gene expression. These findings may indicate that the pattern of IVSC is the reflection of regional differences in plasticity and the result of selective expression of related genes. Finally, our results also highlighted the importance of individual effects when explaining individual differences in cognition.

## Acknowledgements

The authors thank all the volunteers for their participation in the study. This work was supported by the STI2030-Major Projects (2022ZD0213300, 2021ZD0200500), National Natural Science Foundation of China (32271145, 81871425), Fundamental Research Funds for the Central Universities (2017XTCX04), Open Research Fund of the State Key Laboratory of Cognitive Neuroscience and Learning (CNLZD2101, CNLYB2001).

## Online Methods

### Participants

The human MRI data employed in this work came from five datasets, HCP-YA ^[1]^, HCP Retest ^[1]^, HCP-A ^[21]^, Cam-CAN ^[22]^, and BABRI ^[23]^. Each study of the corresponding dataset was approved by the research ethics review boards of the respective institutions. This project was approved by the research ethics review board of Beijing Normal University. These five datasets are described below.

*HCP-YA.* We used the final release of HCP-YA, which includes 1206 healthy young adult participants. We restricted our analysis to subjects from different families to avoid the effects of shared genes and the environment on the evaluation of individual variability (n = 457). After 38 subjects were excluded for low quality in raw diffusion weighted MRI scans or low quality data in the QSIPrep pipeline ^[44]^, the present study included data from a total of 419 healthy young adult participants from the HCP-YA study (221 males, age 22 - 37).

*HCP Retest.* Among the 1206 subjects from the HCP-YA study, 46 subjects underwent two separate scans with an interval ranging from 0.5 to 11 months. Three subjects were excluded after quality control of MRI scans and the QSIPrep pipeline ^[44]^. The two MRI scans of the remaining 43 subjects from HCP Retest were included in this study (13 males, age 22-35).

*HCP-A.* HCP-A is an ongoing study aimed at enrolling 1200+ healthy adults (age 36-100+). We used HCP-A Release 2.0, which included 725 healthy adults. Considering the influence of grey matter atrophy due to ageing on the evaluation of IVSC, we excluded participants who were older than 68 years, which is an upper tertile of ages of all participants from the HCP-A, Cam-CAN and BABRI. After excluding the subjects whose quality in the raw MRI data or QSIPrep pipeline ^[44]^ was low, data from 470 healthy adults from the HCP-A study were included in this study (300 males, age 36-68).

*Cam-CAN.* Cam-CAN enrolled 2681 healthy participants, and 700 of those participants underwent MRI scans. Similarly, participants older than 68 years were excluded. We also excluded participants younger than 30 years because we wanted to focus only on IVSC in the ageing process. Finally, 382 participants from Cam-CAN were included in this study (195 males, age 31-68 years).

*BABRI.* BABRI is an ongoing study based on a community cohort focusing on asymptomatic stages of dementia and aiming to develop prevention strategies for cognitive impairment. Similarly, after excluding participants older than 68 years, 414 healthy participants from the BABRI were included (141 males, age 45-68).

### Cognitive assessment

To explore the association between cognition and individual variability, we only included the cognitive data from the HCP-A and BABRI because both studies used the same neuropsychological tests to assess participants’ executive function and attention. For the HCP-A dataset, in addition to the NIH toolbox, additional cognitive tests, including the Rey auditory verbal learning task, delay discounting, and trials A and B, were administered to all participants as cognitive assessments. For the BABRI dataset, a battery of neuropsychological tests was adopted to assess cognitive function across five domains: memory, executive function, attention, visuospatial ability and language. Among the five domains, executive function was measured via the Trail B, Stroop B and C tests; attention was assessed by the Trail A, symbol digit modalities and Stroop A tests; and memory was examined by the Rey auditory verbal learning task.

### Image acquisition

*HCP-YA and HCP Retest.* MRI data were collected with a customized 3T Connectome Scanner adapted from Siemens Skyra. T1-weighted (T1w) scans used 3D magnetization prepared rapid gradient echo (MPRAGE) (slices = 256, TR = 2400 ms, TE = 2.14 ms, flip angle = 8°, voxel size = 0.7 mm isotropic). A multi-shell diffusion-weighted echo-planar imaging sequence was used for diffusion-weighted MRI (dMRI) data (90 diffusion-weighted directions for b =1000, 2000 and 3000 s/mm^2^ and 18 images with b = 0 s/mm^2^, slices = 111, TR = 5500 ms, TE = 89.50 ms, slice thickness = 1.25 mm, FOV = 210 x 180 mm^2^, and acquisition matrix = 168 x 144).

*HCP-A.* MRI data were acquired with a 3T MRI scanner (Prisma, Siemens). T1w scans used multiecho MPRAGE (sagittal slices = 208, TR = 2500 ms, TE = 1.81/3.6/5.39/7.18 ms, TI = 1000 ms, slice thickness = 0.8 mm, flip angle = 8°, FOV = 256 x 256 mm^2^, and acquisition matrix = 320 x 320). A multi-shell diffusion-weighted echo-planar imaging sequence was used for dMRI data (92 diffusion-weighted directions for b =1500, 93 diffusion-weighted directions for 3000 s/mm^2^, 14 images with b = 0 s/mm^2^, slices = 92, TR = 3230 ms, TE = 89.20 ms, slice thickness = 1.5 mm, FOV = 210 x 210 mm^2^, and acquisition matrix = 140 x 140).

*Cam-CAN.* All MRI datasets were collected at a single site using a 3T MRI scanner (Trio, Siemens). 3D MPRAGE was used for the T1w scan (TR = 2250 ms, TE = 2.99 ms, TI =900 ms, flip angle = 9°, FOV = 256 mm x 240 mm x 192 mm, voxel size = 1 mm isotropic). A twice-refocused spinLecho sequence was used for dMRI (30 diffusion gradient directions for b = 1000 and 2000 s/mm^2^, 3 images with b = 0 s/mm^2^, axial slices = 66, TR = 9100 ms, TE = 104 ms, FOV = 192 mm x 192 mm) *BABRI.* MRI data were acquired with a 3T MRI scanner (Trio, Siemens Magnetom). A sagittal 3D MPRAGE was used for the T1w scan (sagittal slices = 176, TR = 1900 ms, TE = 3.44 ms, TI = 900 ms, slice thickness = 1 mm, flip angle = 9°, FOV = 256 x 256 mm^2^, and acquisition matrix = 256 x 256). A single-shell diffusion-weighted echo-planar imaging sequence was used for dMRI data (30 diffusion-weighted directions with b = 1000 s/mm^2^ and a single image with b = 0 s/mm^2^, axial slices = 70, TR = 9500 ms, TE = 92 ms, slice thickness = 2 mm, FOV = 256 x 256 mm^2^, and acquisition matrix = 128 x 128).

### Image processing

The image processing pipeline consisted of preprocessing and reconstruction workflows. QSIPrep ^[44]^ 0.14.3, which is based on FSL ^[45]^, DSI studio ^[46]^, DIPY ^[47]^, ANTs ^[48]^and MRtrix ^[49]^, is used to perform image processing and create individual white matter networks. Detailed procedures are described below.

*Preprocessing.* For T1w images, ANTs were used to extract the brain data and generate a brain mask. FSL was utilized for brain tissue segmentation. Finally, ANTs were used to normalize the T1w image to MNI space. For dMRI data, images were grouped according to the phase encoding direction. Then, MRtrix was used to denoise diffusion-weighted images. FSL was used for distortion and head motion correction. Finally, the template image with b = 0 was created and registered to the skull-stripped T1w image.

*Reconstruction.* The reconstruction workflow performed advanced reconstruction and tractography methods to create a white matter network. Single-shell and multi-shell constrained spherical deconvolution were separately applied to single-shell and multi-shell dMRI data to estimate voxel-level diffusion models. A probabilistic tractography algorithm in MRtrix was used to produce a fixed number of streamlines, and an anatomically constrained tractography framework was used during tracking. Then, spherical deconvolution-informed filtering of tractograms was applied to improve the quantitative nature of whole-brain streamline reconstructions. Finally, the Brainnetome atlas ^[50]^ was used to define nodes in the white matter network, and the number of streamlines connecting a pair of regions was used to weight edges.

### Analyses of IVSC

#### Harmonization of multi-site data

For removing the site effects in structural connectivity from multiple sites, the ComBat model that was successfully applied to the harmonization of the measures from MRI data, such as cortical thickness ^[24]^ and function connectivity ^[51]^ was utilized. This approach assumes that sites have both additive and multiplicative effects on data. Hence, the model is defined as follows:

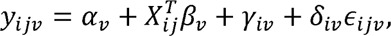

where *α_v_* is the mean value of structural connectivity 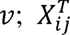 is a design matrix for the covariates *β_v_*; *γ_iv_* and *δ_iv_* are the additive and multiplicative site effects of site; for connectivity *v*, respectively. The final ComBat-harmonized connectivity is then defined as:

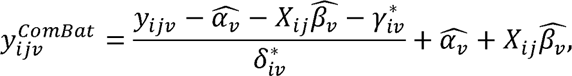

where 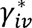 and 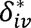 are the estimates of *γ_iv_* and *δ_iv_* using empirical Bayes. In this study, we included age and sex as covariates to preserve biological information in the data.

#### Definition of individual variability of the structural network

The IVSC was quantified according to the definition in previous studies ^[10–12]^. For a given node *i*, the intersubject difference was defined as follows:

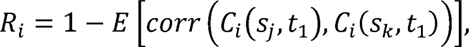

where *j*, *k* = 1,2 … M (j ≠ k); M is the number of participants; *c_i_(s_j_,t_1_*) is a vector of structural connectivity values of node *i* in participant *j* at timepoint *t*_l_.

The intrasubject difference was estimated using the HCP Retest dataset, where each participant underwent two MRI scans. It was defined as follows:

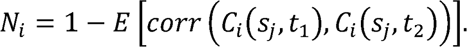

Then, IVSC was defined as follows:

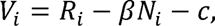

where *β* and *c* are parameters determined via ordinary least-squares in the HCP Retest dataset. These parameters and *N* were applied to other datasets to estimate IVSC.

#### Sliding window analysis

To examine the ageing of structural network variability, cross-subject sliding window analyses were performed for participants from HCP-A, Cam-CAN and BABRI in ascending order of age, with a window length of 10 participants and a step size of 1 participant. Using this method, 1253 overlapping subgroups were generated, and their structural network variabilities and mean ages were calculated.

### Spatial permutation testing (Moran spectral randomization)

To evaluate the association between the spatial distribution of structural network variability and other cerebral properties, a spatial permutation procedure called Moran spectral randomization was used to control the effect of spatial autocorrelation on the association. The inverse Euclidean distance between nodes was used to denote the spatial proximity when using Moran spectral randomization to generate 10000 random brain maps that preserved the spatial autocorrelation of the original map. The null distribution of Spearman’s r values was obtained by relating structural network variability with random brain maps. The p value was computed as the proportion of null r values that were greater than the true r value.

### Transcriptional analyses

#### Transcriptional data preprocessing

Transcriptional data of six post-mortem brains were provided by the Allen Human Brain Atlas ^[52]^. Analyses were limited to the left brain only because only two of six brains included samples from the right hemisphere. The Abagen toolbox ^[53]^ was used to preprocess the transcriptional data according to the suggestions in a previous study^[54]^.

Specifically, microarray probes were reannotated to corresponding genes based on the database provided by Arnatkevičiūtė et al. ^[55]^. Filter noisy probes that did not exceed the background in at least 50% of all samples across all subjects. For each gene, the representative probe with the expression level most correlated to RNA-seq measures collected in two 2 of six brains was selected. Next, tissue samples were assigned to the closest regions in the Brainnetome atlas, while samples more than 2 mm from all regions were not assigned to any regions. To account for interindividual variation, expression values within each brain across regions were normalized with a scaled robust sigmoid function ^[56]^. To retain the genes with consistent expression patterns across six brains, differential stability ^[57]^ was calculated for each gene, and a threshold of 0.1 was imposed. Finally, regional expression values were averaged across donors.

It is important to provide gene specificity on the observed transcriptomic-neuroimaging effect to avoid nonspecific effects ^[58]^. To retain brain-specific genes, genes with higher expression in brain sites than in nonbrain sites were selected using the Genotype-Tissue Expression database (https://www.gtexportal.org/). This procedure produced a matrix with 123 rows corresponding to brain regions and 2112 columns corresponding to the retained genes.

#### PLS analysis

The correlation between gene expression and structural network variability was examined with a PLS analysis that is an unsupervised multivariate method to project two matrices into a lower-dimensional space where the covariance between them is maximal. This analysis was performed using the *PLSRegression* function in the Python package scikit-learn (https://scikit-learn.org/stable/index.html) with the number of components set as 1 because IVSC only has 1 dimension.

#### Bootstrap resampling

To obtain reliable estimation of gene weights, bootstrap resampling was used. The values in the structural network variability vector were randomly selected with replacement 10000 times. PLS analysis was performed using these bootstrapped structural network variability vectors resulting in distributions of gene weights. The reliable gene weights were quantified by the ratio of the true gene weight to the standard error of its bootstrapped distribution. The ratio is the combination of both the value and reliability of gene weights.

#### Gene set enrichment analysis

To further investigate biological pathways related to the genes identified by PLS analysis, the method used by Hansen et al. ^[59]^ was adapted. The latest gene annotation provided by Gene Ontology (geneontology.org) was downloaded on 2^nd^ July 2022. For each biological process in gene annotation, the enrichment score was defined as the mean weights of the genes in the biological process. The null distribution of enrichment scores was constructed by permuting the structural network variability map 10000 times with Moran spectral randomization and repeating PLS analyses and calculating the enrichment score. This analysis was separately performed on the positive genes with 50% of the most positive weights and negative genes with 50% of the most negative weights. For positive genes, significance was measured by the proportion of null enrichment scores that were greater than the true enrichment score. For negative genes, the proportion of null enrichment scores that were smaller than the true enrichment score was utilized to quantify significance.

To determine cell types where the genes identified by PLS are preferentially expressed, the method performed by Hansen et al. ^[59]^ was adopted. Briefly, gene annotation about cell type was provided by Seidlitz et al. ^[60]^, who used hierarchical clustering based on spatial expression profiles to distinguish 7 canonical cell types: astrocytes, endothelial cells, microglia, excitatory neurons, inhibitory neurons, oligodendrocytes and oligodendrocyte precursors. For each cell type, the enrichment score was defined as the proportion of genes that were identified by PLS analysis and preferentially expressed in that type as well. This analysis was separately performed on the positive genes with 50% of the most positive weights and negative genes with 50% of the most negative weights. A nonparametric permutation test was used to quantify the significance.

#### Development of gene expression

We used BrainSpan ^[61]^, a database of gene expression across development, to depict the development of the gene expression component determined by PLS analysis in the Allen Human Brain Atlas. Post-mortem brains in BrainSpan were grouped into 5 developmental stages: foetus (8–37 postconception week), infant (4 month – 1 year), child (2–8 year), adolescent (11–19 year) and adult (21–40 year). For each group, gene expression scores in 16 regions were calculated by multiplying the gene weights derived from the PLS analysis in the Allen Human Brain Atlas and mean expression across post-mortem brains in the group.

To relate gene expression scores in BrainSpan and structural network variability, we mapped the Brainnetome Atlas into 16 Brodmann areas that are included in BrainSpan. For each Brodmann area, structural network variability was averaged across the Brainnetome Atlas regions corresponding to the Brodmann area. Then, the relationship between gene expression scores and structural network variability was examined in each developmental stage.

### Cognitive prediction

#### Modified ridge regression

To test whether the structural connectivity involved in cognitive tasks differed across individuals, we used two models to predict individual cognition. One was ridge regression, which is formulized as follows:

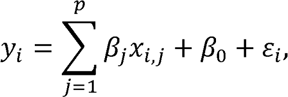

where *Y_i_* is the cognitive score of individual *i*, *x_i,j_* is the weight of structural connectivity *j* of individual *i*, *β_j_* is the regression coefficient and *ε_i_* is the error. Ridge regression has been shown to have similar performance to other machine learning methods but is computationally more efficient when using high-dimensional imaging data as features ^[7]^. When fitting the model, ridge regression used an L2 penalty to avoid overfitting and to improve generalizability. The objective function is:

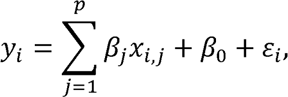

where *α* is a hyperparameter that controls the penalty of model complexity.

Considering that the structural connectivity encoding cognitions may be different across individuals, we proposed a modified ridge regression that is a neural network with a hidden layer. The model can be defined as follows:

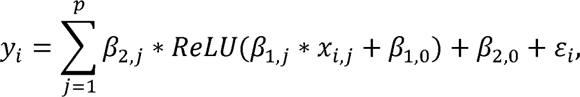

where *β*_1,*<u>j</u>*_ is the coefficient to linearly transform structural connectivity, *β*_2,*j*_ is the coefficient to predict cognition and *ReLU*(·) is a linear rectification function. *ReLU*(·) and *β*_1,*j*_ together determine which features are used to predict cognition.

Similarly, the modified ridge regression’s objective function is:

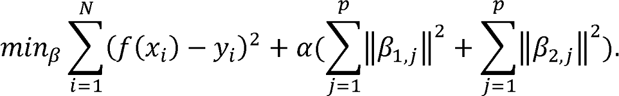

#### Prediction framework

To train models of cognitive prediction, a nested 2F-CV where the outer 2F-CV was used in the evaluation of models’ performance and the inner 2F-CV was used to determine the optimal hyperparameter *α*.

*Outer 2F-CV.* For the outer 2F-CV, we merged the HCP-A and BABRI datasets because they both used Trail B and Trail A to measure individuals’ executive function and attention, respectively. Then, participants in the merged dataset were sorted according to the outcome (i.e., executive function or attention). The participants with an odd rank were assigned to subset 1, and other participants were assigned to subset 2. In the first fold, subset 1 was used as the training dataset, and subset 2 was used as the test dataset. The second fold was the opposite. Each feature was normalized across the training dataset, and the mean value and standard deviation derived from the training dataset were also applied to the test dataset. After the optimal hyperparameter *α* was selected, a model with the optimal *α* was trained using all participants in the training dataset and then applied to the test dataset to predict the outcome. Pearson’s correlation coefficients and MAE between the predicted and true outcomes of participants in the test dataset were calculated to quantify the prediction accuracy.

*Inner 2F-CV.* For the inner 2F-CV, the training dataset was further split into 2 subsets like for the outer 2F-CV. These subsets were used to train models under different values of hyperparameter *α* ([2^-4^, 2^-3^, …, 2^4^, 2^5^]) in turn, and these models were applied to the remaining dataset to calculate Pearson’s correlation coefficient and MAE between the predicted and true outcomes. For each value of *α*, Pearson’s correlation coefficients and MAE were averaged across 2 inner loops. Then, the sum of the normalized mean Pearson’s correlation coefficient and normalized reciprocal of the mean MAE was defined as the inner prediction accuracy. The value of hyperparameter *α* with the highest inner prediction accuracy was selected as the optimal *α*.

*Randomly split 2F-CV.* To validate whether the result is influenced by data splitting, we randomly split the merged dataset into two halves in both outer 2F-CV and inner 2F-CV, and calculated the mean Pearson’s correlation coefficient and MAE across two outer folds. This procedure was repeated 100 times to obtain the distribution of mean Pearson’s correlation coefficients and MAE.

*Significance of prediction accuracy*. A permutation test was performed to evaluate the significance of the prediction accuracy. Briefly, we performed nested 2F-CV 1000 times with permuting outcomes across the training dataset in each run. Null distributions of Pearson’s correlation coefficients and MAE were obtained. The significance of Pearson’s correlation coefficient was determined as the proportion of the null coefficient higher than the true coefficient, and the significance of MAE was defined as the proportion of null values lower than the true value.

### Replication analyses

#### Effects of edge threshold

To evaluate whether the IVSC pattern identified in this study is robust to edge threshold, we used different edge thresholds to remove weak edges in structural networks and then re-calculated the IVSC maps. The Spearman’s correlations between the IVSC map without removing edges and the IVSC maps with removing weak edges were used to assess the stability of IVSC pattern.

#### Effects of brain parcellation

In addition to the Brainnetome atlas, we also used AAL and Gordon’s atlases to define network nodes. The IVSC maps were separately calculated from the networks based on AAL and Gordon’s atlases and analysed with the similar procedure that was performed in the networks based on Brainnetome atlas.

## Data availability

HCP-YA, HCP Retest and HCP-A are available at https://www.humanconnectome.org/. Cam-CAN is available at https://www.cam-can.org/index.php?content=dataset. BABRI is available upon on a request. Allen Human Brain Atlas available at https://help.brain-map.org/display/humanbrain/Allen+Human+Brain+Atlas. The Genotype Tissue Expression database is available at https://www.gtexportal.org/.

## Code availability

The code for generating individual variability in structural connectivity, statistics and cognitive prediction is available on GitHub (https://github.com/forwho/Individual-Difference-in-Structural-connectivity).

