## Supplemental figure for "Individual Variability in the Structural Connectivity Architecture of the Human Brain"

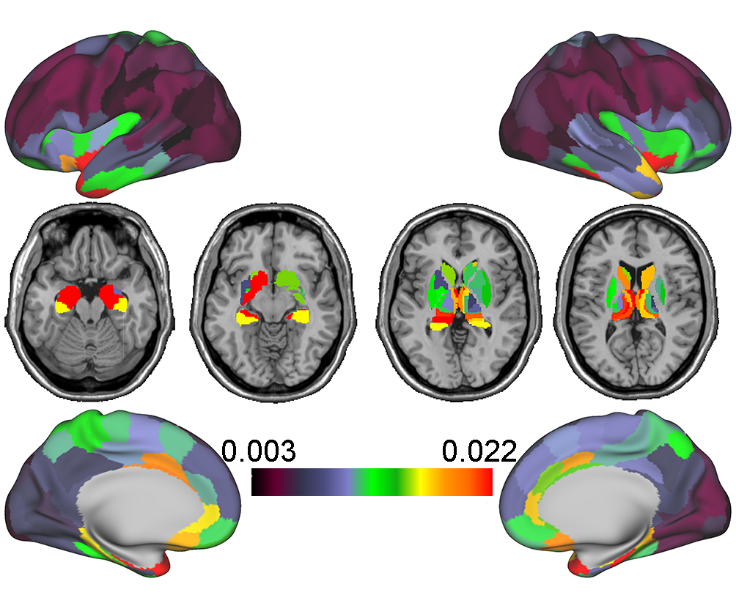

**Figure S1. Intraindividual differences in structural connectivity.** The intraindividual differences in structural connectivity were calculated based on HCP Rest datasets. HCP, Human Connectome Project.

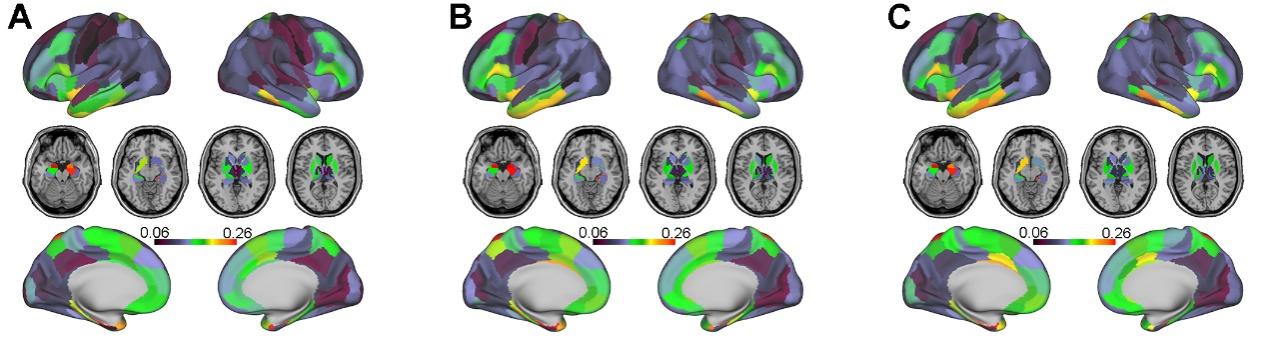

**Figure S2. Individual variability in structural connectivity maps derived from three different datasets. A,** Individual variability in structural connectivity maps derived from the HCP-A. **B,** Individual variability in structural connectivity maps derived from the Cam-CAN. **C,** Individual variability in structural connectivity maps derived from the BABRI. HCP-A, Human Connectome Project Ageing, Cam-CAN, Cambridge Centre for Ageing and Neuroscience; BABRI, Beijing Ageing Brain Rejuvenation Initiative;

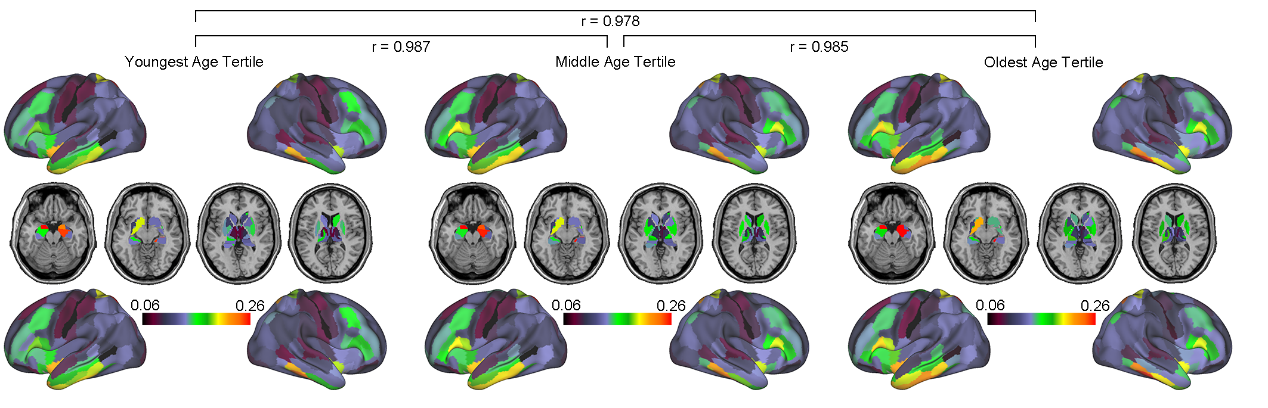

**Figure S3. Individual variability in structural connectivity in three age-based tertiles.** We split the samples in the combined dataset into age tertiles and calculated individual variability in structural connectivity in each group. We found that individual variability in structural connectivity was nearly identical in each of the three age-based tertiles.

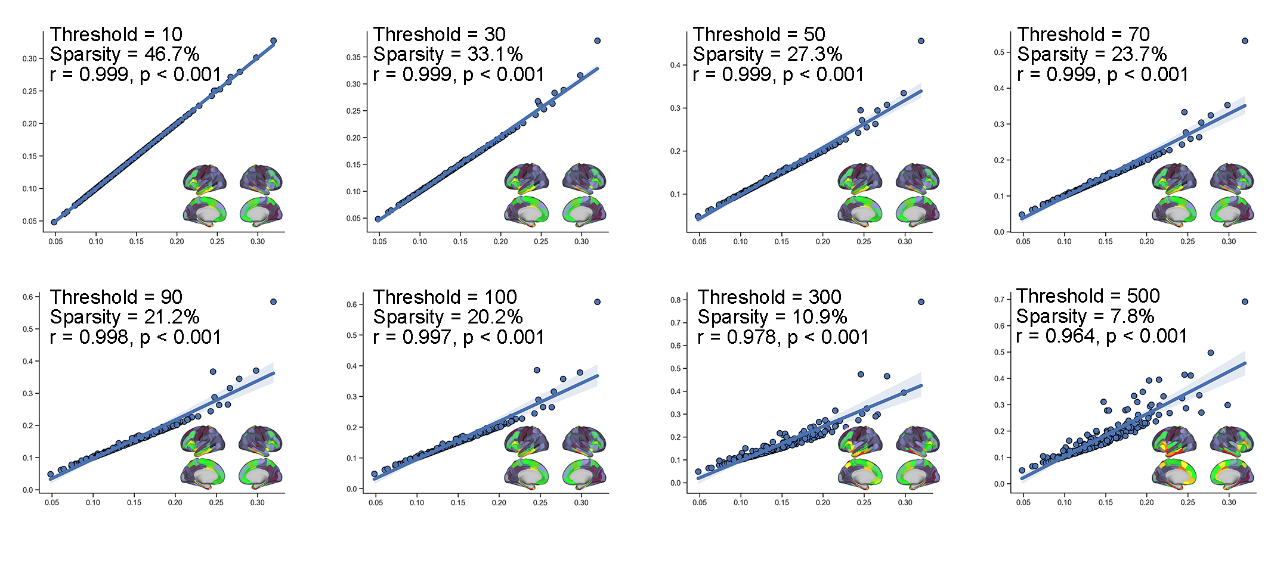

**Figure S4. Effects of edge thresholds on IVSC measurement.** We used different edge thresholds to remove weak edges in structural networks and then re-calculated the IVSC maps. We found that the IVSC maps after removing weak edges were highly similar to the IVSC map without removing edges. Inset plots show the IVSC maps with removing weak edges under different threshold. IVSC, individual variability in structural connectivity.

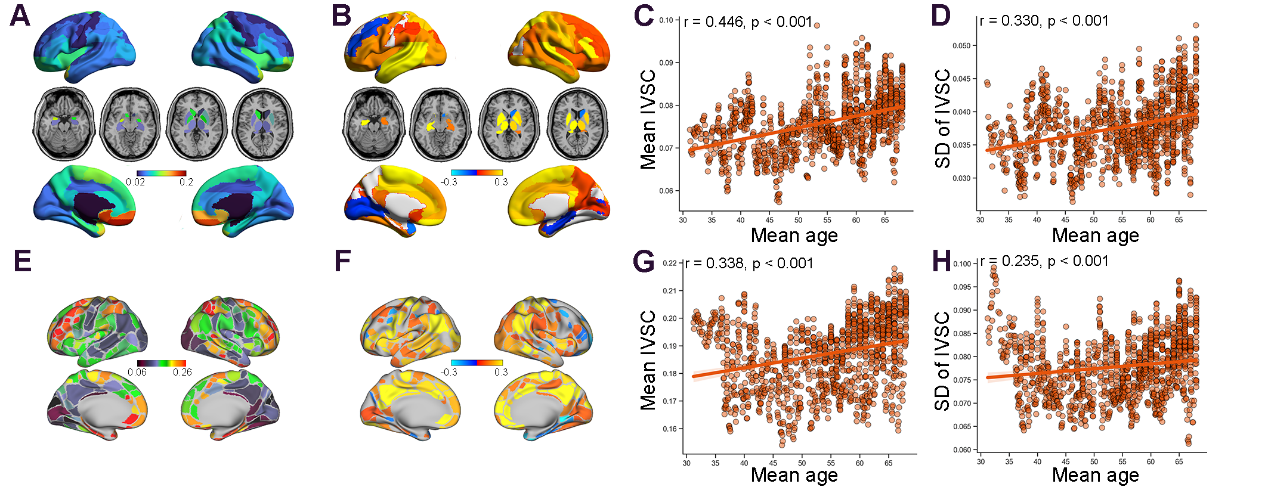

**Figure S5. Temporal spatial pattern of IVSC based on AAL and Gordon’s atlases. A,** The IVSC map based on AAL atlas. **B,** Age-related rates of IVSC based on AAL atlas. **C,** the relationship between mean IVSC based on AAL atlas and mean age. **D,** The relationship between the standard deviation of IVSC based on AAL atlas and mean age. **E,** The IVSC map based on Gordon’s atlas. **F,** Age-related rates of IVSC based on Gordon’s atlas. **G,** the relationship between mean IVSC based on Gordon’s atlas and mean age. **H,** The relationship between the standard deviation of IVSC based on Gordon’s atlas and mean age. IVSC, individual variability in structural connectivity; AAL, automated anatomical labelling.

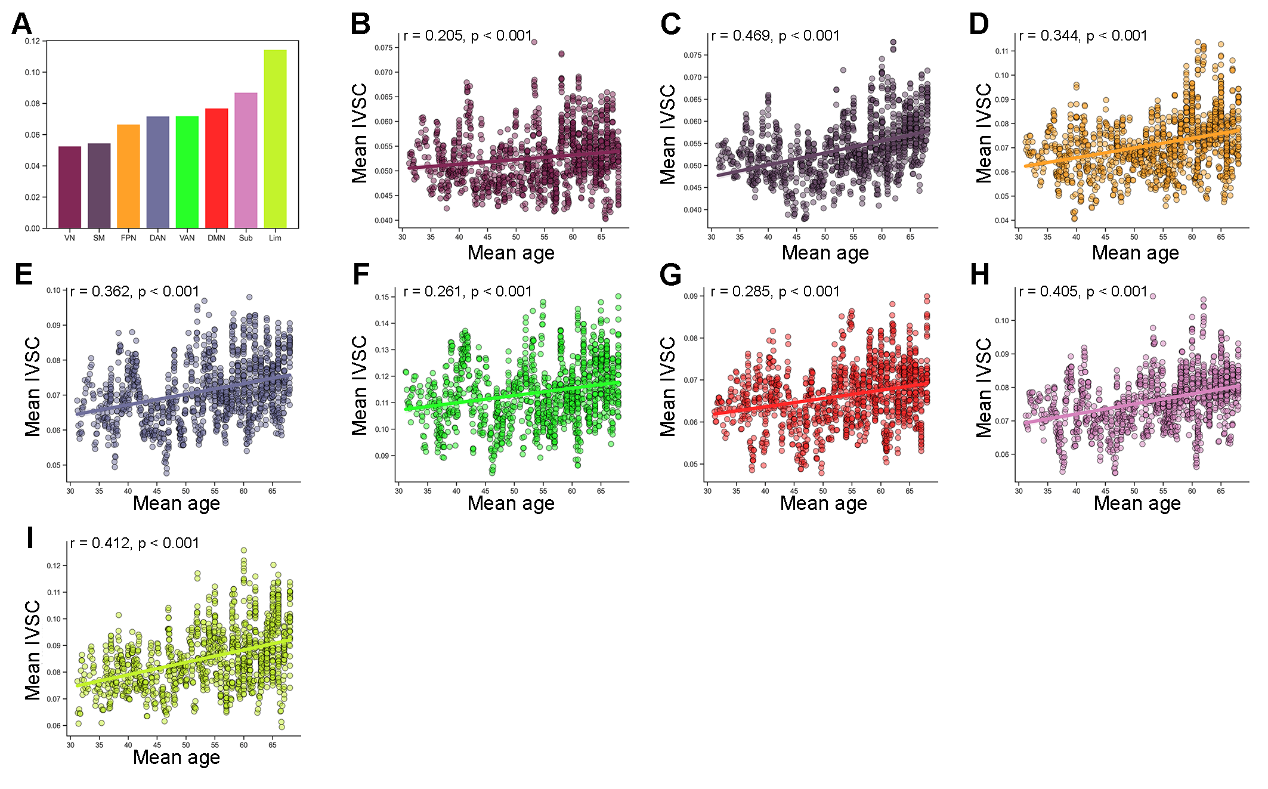

**Figure S6. Quantify the IVSC map based on AAL atlas in 8 brain systems. A,** Mean IVSC in 8 brain systems. **B-I,** The relationship between mean IVSC in 8 brain systems and mean age. AAL, automated anatomical labelling; IVSC, individual variability in structural connectivity; SM, somatomotor network; VN, visual network; DMN, default mode network; DAN, dorsal attention network; Sub, subcortical system; FPN, frontosparietal network; VAN, ventral attention network; Lim, limbic network.

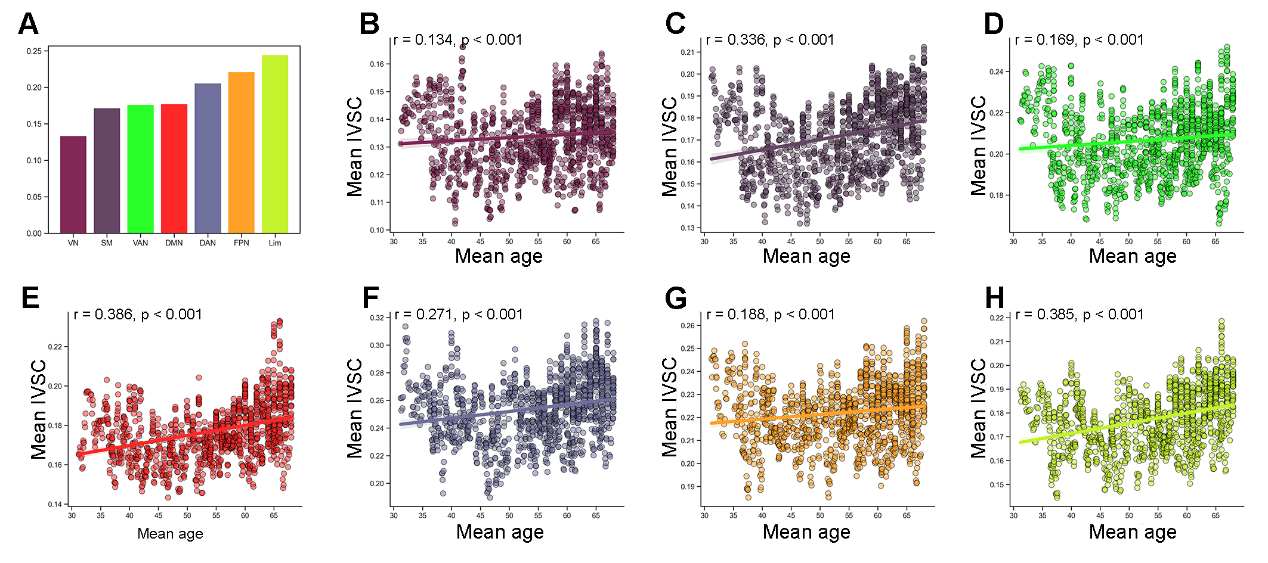

**Figure S7. Quantify the IVSC map based on Gordon’s atlas in 8 brain systems. A,** Mean IVSC in 8 brain systems. **B-H,** The relationship between mean IVSC in 8 brain systems and mean age. IVSC, individual variability in structural connectivity; SM, somatomotor network; VN, visual network; DMN, default mode network; DAN, dorsal attention network; FPN, frontosparietal network; VAN, ventral attention network; Lim, limbic network.

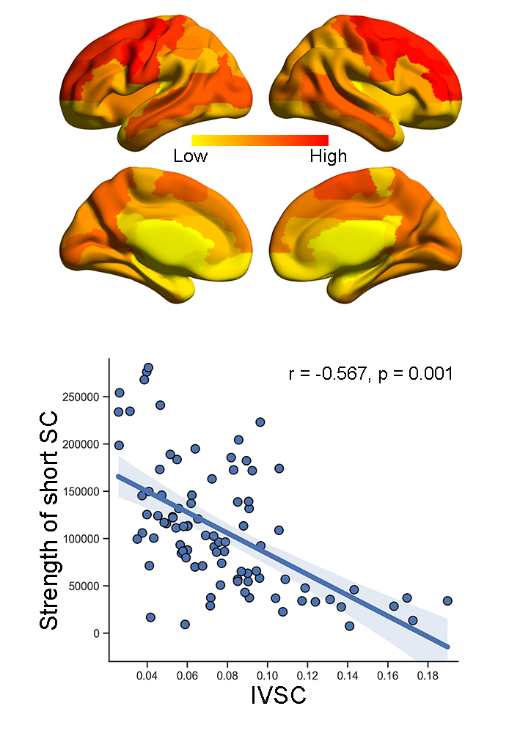

**Figure S8. The relationship between IVSC based on AAL atlas and the strength of short SC.** IVSC, individual variability in structural connectivity; AAL, automated anatomical labelling.

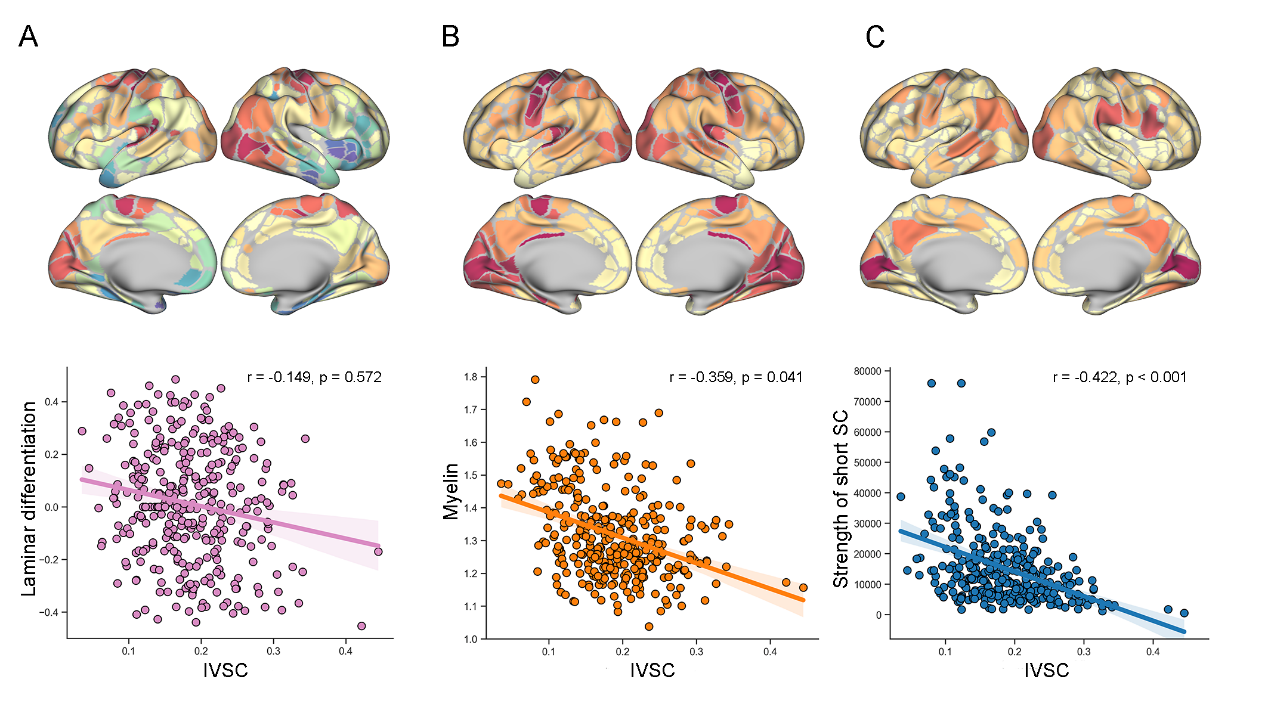

**Figure S9. The relationship between IVSC based on Gordon’s atlas and fundamental properties of brain organization. A,** Spatial distribution of the laminar differentiation identified by Paquola et al [62]. **B,** Spatial distribution of myelin content measured by T1w/T2w mapping [63]. **C,** Spatial distribution of short connectivity strength. **D-F,** The relationship between IVSC with laminar differentiation, myelin content and strength of short connectivity. IVSC, individual variability in structural connectivity; SC, structural connectivity. IVSC, individual variability in structural connectivity; SC, structural connectivity.

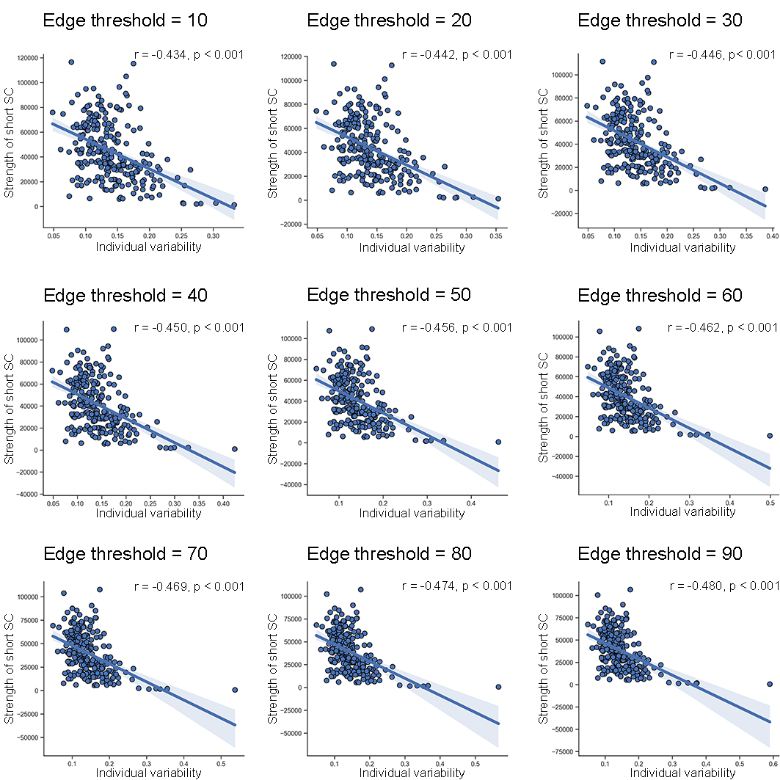

**Figure S10. The effect of edge thresholds on the correlation between individual variability in SC and strength of short SC across brain regions.** To evaluate whether the correlation was stable across different edge thresholds. We recalculated individual variability in SC after filtering the weak edges whose weights were smaller than a threshold and then examined the correlation between individual variability in SC and the strength of short SC. We found that the relationship was stable across different edge thresholds. SC, structural connectivity.

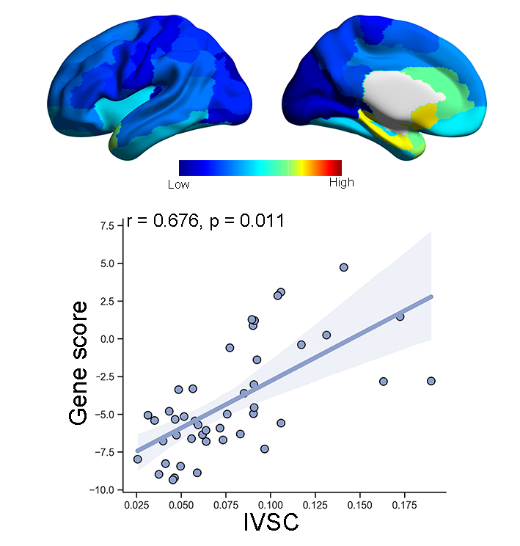

**Figure S11. The relationship between IVSC based on AAL atlas and gene expression.** IVSC, individual variability in structural connectivity; AAL, automated anatomical labelling.

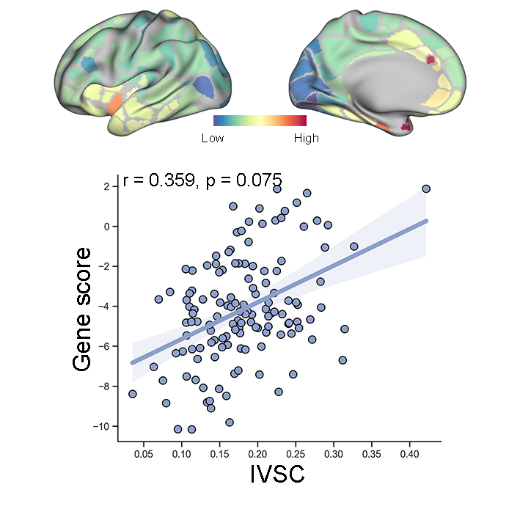

**Figure S12. The relationship between IVSC based on Gordon’s atlas and gene expression.** IVSC, individual variability in structural connectivity; AAL, automated anatomical labelling.
