## Supplementary material for "Individual Variability in the Structural Connectivity Architecture of the Human Brain": Table 1

**Table 1. Demographical information of participants from 5 datasets.**

| Dataset | Sex (Male/Female) | Age (Year) | Trails A | Trails B |
| --- | --- | --- | --- | --- |
| HCP Retest | 13/30 | 22.0 - 35.0 (30.3 ± 3.3) | - | - |
| HCP-YA | 221/198 | 22.0 - 37.0 (28.3 ± 4.0) | - | - |
| HCP-A | 200/270 | 36.0 - 67.9 (51.0 ± 9.1) | 1.2 - 317.7 (66.6 ± 36.9) | 0.3 -78.5  (26.9 ± 10.3) |
| Cam-CAN | 195/185 | 31.0 - 68.0 (49.6±11.1) | - | - |
| BABRI | 141/271 | 45.0 - 68.0 (61.6 ± 4.5) | 60.0 - 318.0 (146.2 ± 45.9) | 24.0 - 115.0 (53.0 ± 14.0) |

HCP-YA, Human Connectome Project Young Adults; HCP-A, Human Connectome Project Ageing; Cam-CAN, Cambridge Centre for Ageing and Neuroscience; BABRI, Beijing Ageing Brain Rejuvenation Initiative.
